# Defining chromatin state transitions predicts a network that modulates cell wall remodeling in phosphate-starved rice shoots

**DOI:** 10.1101/706507

**Authors:** Maryam Foroozani, Sara Zahraeifard, Dong-Ha Oh, Guannan Wang, Maheshi Dassanayake, Aaron Smith

**Author notes:** Author contributions: M.F. performed the experiments, analyzed the data, and wrote the article; S.Z provided technical assistance; D-H.O. and G.W. provided bioinformatics assistance; M.D. supervised the data analysis; A.S. conceived the project, supervised the experiments and data analysis, complemented the writing, and agrees to serve as the author responsible for contact and ensure communication. Funding information: Funding was provided by a Plant Genome Research Program grant from the National Science Foundation (IOS-1127051). Author for contact: A. Smith.

## Abstract

Phosphorus (P) is an essential plant macronutrient vital to fundamental metabolic processes. Plant-available P is low in most soils, making it a frequent limiter of growth. Declining P reserves for fertilizer production exasperates this agricultural challenge. Plants modulate complex responses to fluctuating P levels via global transcriptional regulatory networks. Although chromatin structure plays a substantial role in controlling gene expression, the chromatin dynamics involved in regulating P homeostasis have not been determined. Here we define distinct chromatin states across the rice genome by integrating multiple aspects of chromatin structure, including the H2A.Z histone variant, H3K4me3 modification, and nucleosome positioning. In response to P starvation, 40% of all protein-coding genes exhibit a transition from one chromatin state to another at their transcription start site. Several of these transitions are enriched in subsets of genes differentially expressed by P deficiency. The most prominent subset supports the presence of a coordinated signaling network that targets cell wall structure and is regulated in part via a decrease of H3K4me3 at the transcription start site. The P-starvation induced chromatin dynamics and correlated genes identified here will aid in enhancing P-use efficiency in crop plants, benefitting global agriculture.

**One sentence summary:** Combining data for three components of chromatin structure from control and phosphate-starved rice shoots reveals specific chromatin state transitions that correlate with subsets of functionally distinct differentially-expressed genes.

## Introduction

Phosphorus (P) is among the most limiting essential nutrients for plants. This is because the primary plant-available form of P, inorganic phosphate (Pi), has poor solubility in most soils (Holford, 1997). As a result, P fertilization of soils is required for crop plants to achieve adequate yields. Unfortunately, P fertilization can result in serious environmental concerns due to nutrient run-off, which will worsen in the future due to the non-renewable nature of P resources (Vance et al., 2003). It is, therefore, necessary to investigate the underlying mechanisms involved in regulating P homeostasis, so as to increase the efficiency of plants to acquire and recycle P. In order to tolerate low-Pi conditions and maintain optimal P levels, plants have evolved a number of physiological, morphological and biochemical responses, such as reduced growth, altered root system architecture, and secretion of organic acids, phosphatases, and nucleases to acquire more Pi (Secco et al., 2013). These responses are modulated by large transcriptional networks in which the MYB protein PHR1 and related transcription factors play key roles (Secco et al., 2013, Sun et al., 2016).

In eukaryotic cells, genes are complexed with core histones and other chromosomal proteins in the form of chromatin. The basic repeating unit of chromatin, the nucleosome, is composed of two copies of each of the four core histones H2A, H2B, H3, and H4 wrapped by 146 bp of DNA (Luger et al., 1997). Therefore, chromatin structure is a key determinant of gene expression. Despite the fact that a large transcriptional cascade governs responses to low-Pi, relatively little is known regarding the associated chromatin dynamics, although evidence for chromatin-level mechanisms modulating Pi deficiency responses is emerging. Smith et al. (2010) demonstrated that mutation of the actin-related protein (ARP) gene, *ARP6*, which is a key component of the SWR1 complex that catalyzes H2A.Z deposition (Deal et al., 2007), resulted in decreased H2A.Z localization at a number of Pi deficiency response genes that were also de-repressed. These changes in H2A.Z and expression also occurred in Pi-deficient wild-type plants (Smith et al., 2010). Recently, we demonstrated a similar phenomenon in rice in which genome-wide H2A.Z distribution was altered similarly by Pi starvation or RNAi knock-down of *ARP6* (Zahraeifard et al., 2018). We also showed that deposition of rice H2A.Z in gene bodies largely resulted in down-regulation, whereas H2A.Z at the TSS was positively or negatively correlated with gene expression, depending on the particular Pi deficiency response genes affected. In a separate study we revealed that changes in nucleosome occupancy correlated with genes differentially expressed by Pi starvation, implicating nuclesome remodelers in modulating Pi deficiency responses in rice (Zhang et al., 2018). Finally, two chromatin-related components have been shown to play roles in Pi-deficiency induced root hair growth in Arabidopsis. The *ALFIN-LIKE 6* (*AL6*) gene encodes a plant homeodomain (PHD)-containing protein that recognizes H3K4 trimethylation and appears to promote enhanced root hair growth during low-Pi conditions by targeting H3K4me3-marked target genes, such as *ETC1*, which functions in root hair cell patterning (Chandrika et al., 2013). The second factor necessary for normal induction of root hair growth in response to Pi deficiency is Arabidopsis *HDA19*, which encodes a histone deacetylase necessary for low-Pi root hair elongation through its role in regulating epidermal cell length (Chen et al., 2015).

Many mechanisms exist to alter the structural characteristics of chromatin, including positioning of nucleosomes, the presence of histone variants, and post-translational modifications of histones (Mariño-Ramírez et al., 2005, Venkatesh and Workman, 2015). Defining the patterns, or states, of chromatin structure by examining multiple marks simultaneously in their spatial context is more informative to understanding transcriptional changes in response to stress (Ernst and Kellis, 2012, Pan et al., 2017). This is exemplified by two recent studies in rice that defined distinct chromatin states by combining multiple histone marks and showed various associations between particular chromatin states and genes differentially expressed by ionizing radiation (Pan et al., 2017) or salinity stress (Zheng et al., 2019). In contrast, no studies have defined chromatin state transitions linked to Pi deficiency responses in plants. Herein we characterized the impact of Pi starvation on the major histone mark, H3K4me3, as well as on chromatin states generated from the combined occupancy data of H3K4me3, H2A.Z, and nucleosomes. The data reveal several distinct chromatin state transitions that accompany expression changes of key subsets of Pi starvation-response genes.

## Results

### H3K4me3 is prominent at the 5’ end of rice protein-coding genes and co-localizes with the H2A.Z histone variant

Previously we demonstrated that dynamics of nucleosome occupancy (Zhang et al., 2018) and H2A.Z deposition (Zahraeifard et al., 2018) were linked to genes differentially expressed in response to phosphate (Pi) starvation in rice shoots. The primary goal herein was to evaluate the combined role of nucleosome occupancy, H2A.Z and another major determinant of chromatin structure, H3K4me3, in modulating responses to Pi starvation. We began by determining the genome distribution of H3K4me3 via ChIP-seq on shoots from 36-day rice (*Oryza sativa* ssp. japonica cv. Nipponbare) seedlings (Supplemental Table S1). Genes were categorized into four groups based on the MSU7 genome annotation: protein-coding genes (PCG), ‘pseudogenes’ (PSG, i.e. annotated genes that are neither expressed nor transposable element-related), transposable element-related genes that are expressed (TEG), and transposable element-related genes not expressed (TE) (Kawahara et al., 2013, Zhang et al., 2018). A prominent H3K4me3 peak was present immediately downstream of the transcription start sites (TSS) of PCG (Figure 1A), similar to previous studies (Zhang et al., 2009, Van Dijk et al., 2010, Du et al., 2013, Zong et al., 2013). In contrast to PCG, H3K4me3 abundance was relatively low at PSG, TEG, and TE (Figure 1A). Next we examined whether sub-groups of PCG exhibited different H3K4me3 patterns. Sorting all PCG according to size revealed a strong correlation between H3K4me3 and gene length (Supplemental Figure S1A,B), indicating that the general pattern of H3K4me3 among all PCG is relatively consistent (i.e. a major peak of H3K4me3 at the TSS). Although a TSS-localized peak of H3K4me3 was observed at virtually all PCG, the abundance of the peak varied. Clustering analysis at a 100-bp window across the TSS revealed 4 distinct clusters of H3K4me3 abundance (Supplemental Figure S1C,D). Gene Ontology (GO) enrichment analysis showed that the clusters with high and moderate abundance were enriched (FDR<0.05) with housekeeping genes, whereas the clusters with relatively low H3K4me3 abundance were enriched in stress-responsive genes (Supplemental Dataset 1).

**Figure 1.**
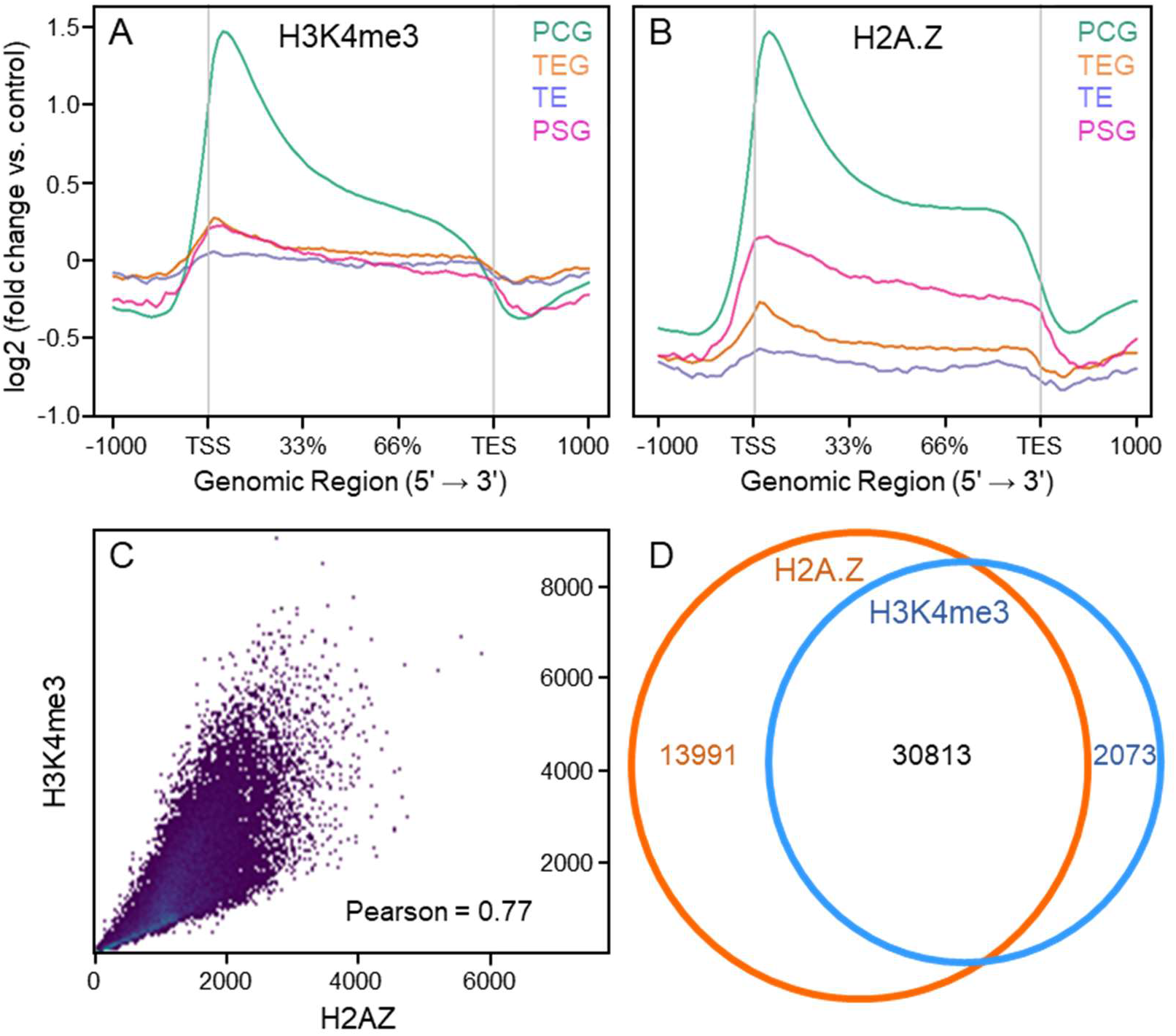
H3K4me3 abundance is predominantly associated with the transcription start site (TSS) and co-localizes with H2A.Z. Distribution of H3K4me3 (A) and H2A.Z (B) among four gene types in shoots from rice seedlings grown under control conditions. Control input reads were used for ChIP-Seq read normalization. PCG, protein coding genes; PG, pseudogenes; TE, non-expressed transposable element-related genes; TEG, expressed transposable element-related genes. (C) Scatter plot of read counts from H3K4me3 and H2A.Z samples (Pearson correlation = 0.77). (D) Venn diagram showing the number of H3K4me3- and H2A.Z-enrichment peaks and the overlap.

The H3K4me3 localization at different gene types (Figure 1A) are similar to those we recently demonstrated for the H2A.Z histone variant (Zahraeifard et al. 2018; Figure 1B). A key difference is that abundance of H3K4me3 is relatively higher than H2A.Z across TEG and TE. To further examine the association between H3K4me3 and H2A.Z we first computed a correlation coefficient using deepTools (Ramírez et al., 2016), which showed that both chromatin marks were correlated across the rice genome (r = 0.77; Pearson correlation coefficient; Figure 1C). Next we identified and compared distinct H3K4me3 and H2A.Z peaks using SICER (Zang et al., 2009) and BEDTools (Quinlan and Hall, 2010). This identified 32,886 H3K4me3 peaks and 44,804 H2A.Z peaks, of which 30,813 (93% of H3K4me3 peaks) overlapped (Figure 1D), showing substantial co-localization of these chromatin marks.

### H3K4me3 and H2A.Z abundance have distinct correlations with gene expression in rice

To compare H3K4me3 abundance with gene expression, we analyzed our previously obtained RNA-seq data (Zahraeifard et al., 2018) from shoot tissues of 36-day rice seedlings (Supplemental Table S1). PCG were ranked according to FPKM and divided into five expression quintiles, as well as a sixth group of genes that were not expressed (i.e. FPKM = 0). We found a clear, positive correlation between transcript abundance and H3K4me3 localization around the TSS (Figure 2A,B), consistent with studies from a variety of species (Bernstein et al., 2002, Santos-Rosa et al., 2002, Barski et al., 2007, Zhang et al., 2009, Van Dijk et al., 2010). In contrast, transcript abundance exhibited a general negative correlation with TES- and gene body-localized H3K4me3 (Figure 2A,B). Genes exhibiting no expression were severely depleted in H3K4me3 at the TSS, but had a moderate level of gene-body H3K4me3. Next, we compared the correlation between H3K4me3 abundance and gene expression with that of H2A.Z (Zahraeifard et al., 2018)(Figure 2C,D). As with H3K4me3, non-expressed genes were deficient in H2A.Z at the TSS, but contained moderate levels of gene-body H2A.Z. However, for the expression quintiles, H2A.Z exhibited a general negative correlation with expression at both the TSS and, especially, in the gene body. Together these results show distinct and overlapping genic patterns of H3K4me3 and H2A.Z, and suggest that the ratio of the two marks is important in the modulation of gene expression.

**Figure 2.**
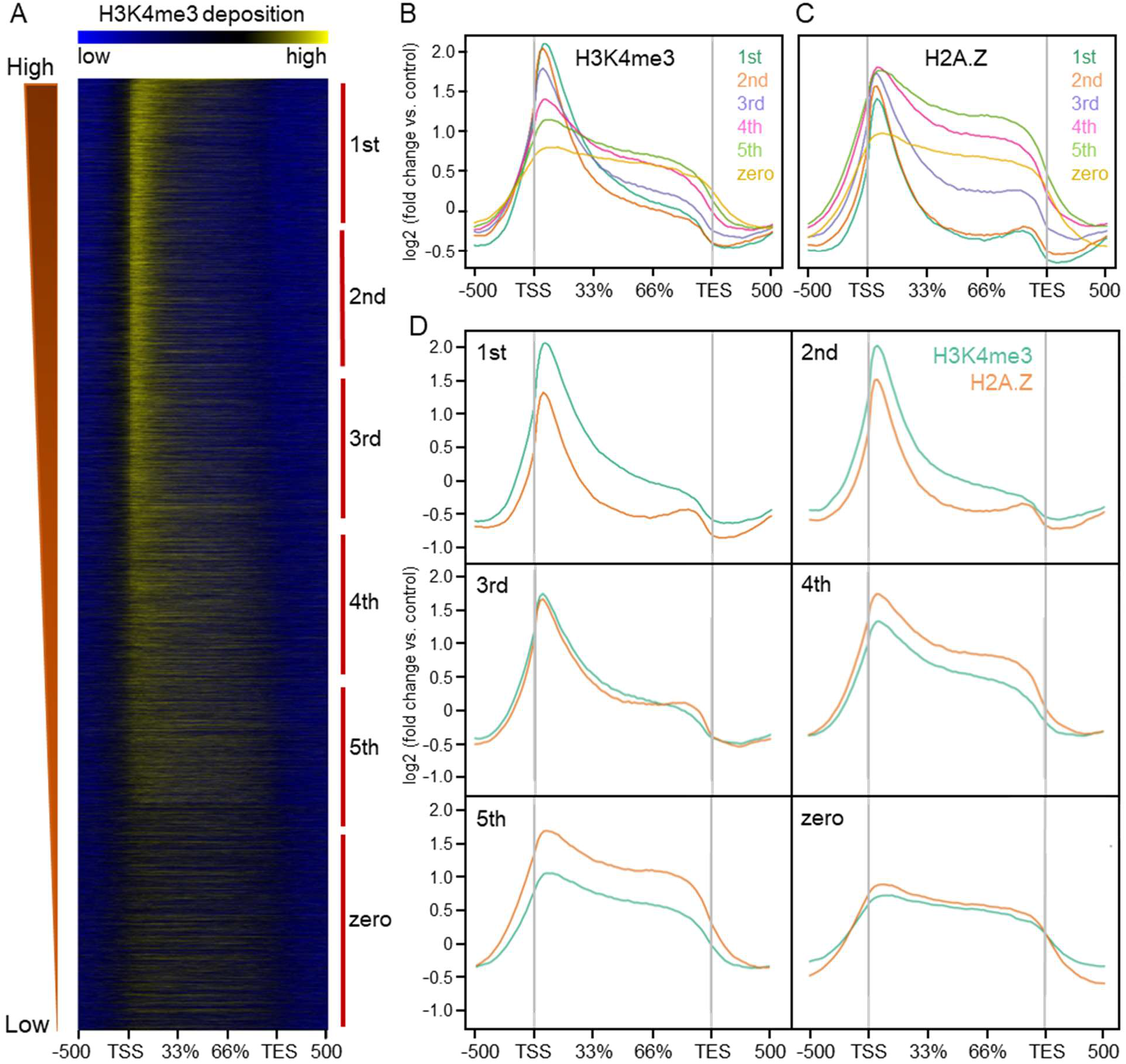
Correlations between H3K4me3 and H2A.Z distribution and gene expression in rice shoots. Heat map (A) and distribution of H3K4me3 (B) across the gene body in control samples for six gene groups ordered based on transcript abundance level (FPKM), defined as 1st (highest) to 5th (lowest) and no expression (zero). (C) Distribution of H3K4me3 and H2AZ across the gene body from 500 bp upstream of the TSS to 500 bp downstream of the TES. Control input reads were used for ChIP-Seq read normalization.

### Pi-starvation impacts H3K4me3 localization

To evaluate a potential role for H3K4me3 in modulating Pi-deficiency responses, we carried out H3K4me3 ChIP-seq on shoots from plants subjected to a 24-hour Pi-deficiency treatment (Supplemental Table S1). As shown in Supplemental Figure S2, Pi-deficiency altered H3K4me3 distribution at PCG, such that the prominent 5’ peak was reduced. These data along with our prior studies (Zahraeifard et al., 2018, Zhang et al., 2018) indicate that nucleosome occupancy, H2A.Z, and H3K4me3 each exhibit distinct changes in response to Pi starvation.

### H3K4me3, H2A.Z, and nucleosome occupancy define five chromatin states in the rice genome

It is becoming increasingly clear that examining multiple chromatin marks simultaneously provides a more robust picture of the dynamic chromatin structure linked to various developmental processes and responses to stimuli (Pan et al., 2017, Yan et al., 2019). Therefore, we integrated our H3K4me3 ChIP-Seq, H2A.Z ChIP-Seq (Zahraeifard et al., 2018) and MNase-Seq data sets (Zhang et al., 2018) to define distinct chromatin states using ChromHMM (Ernst and Kellis, 2012). ChromHMM employs a multivariate Hidden Markov Model that scores the presence or absence of each chromatin mark to determine the major recurring combinatorial and spatial patterns of marks, i.e. chromatin states. ChromHMM identified five chromatin states (CS), each distinguishable from the others by differential enrichment of one or more of the marks tested (Figure 3A). CS1 and CS2 were each deficient in both H2A.Z and H3K4me3, CS3 was enriched in only H2A.Z, CS4 was enriched in both H2A.Z and H3K4me3, and CS5 was enriched in only H3K4me3. Regarding nucleosome density, CS2 and CS3 had moderately higher nucleosome enrichment compared to the other 3 states. Next we mapped the distribution of the five chromatin states across the genome, which revealed biases with a number of genomic features (Figure 3B,D). CS1 was the major chromatin state, accounting for 63 *%* of the rice genome, and was enriched at TE and TEG. It should be noted that highly repetitive regions of the genome were likely designated CS1 due to low numbers of mappable reads rather than bona fide depletion of the chromatin marks examined. TE and TEG were also enriched in CS2 and CS5. This means that the transposable element-related loci were either deficient in both H2A.Z and H3K4me3 or contained H3K4me3 only. In contrast, PSG were enriched in CS2 and CS3, consistent with depletion of both H2A.Z and H3K4me3 or enrichment of only H2A.Z. Finally, PCG were enriched in CS4, consistent with enrichment of both H3K4me3 and H2A.Z. To more specifically characterize PCG, we calculated enrichments at the TSS, TES, and 1kb regions that encompass the TSS or TES (TSS 1kb region: 200 bp upstream to 800 bp downstream of the TSS; TES 1kb region: 800 bp upstream to 200 bp downstream of the TES; Figure 3B). Compared to all bins within PCG, the TSS was more enriched in CS4, CS5, and CS3, whereas the TES was more enriched in CS3 and less enriched in CS4. These results indicate for PCGs generally an overall high occupancy of H2A.Z and/or H3K4me3 at the TSS, but an enrichment of only H2A.Z at the TES.

**Figure 3.**
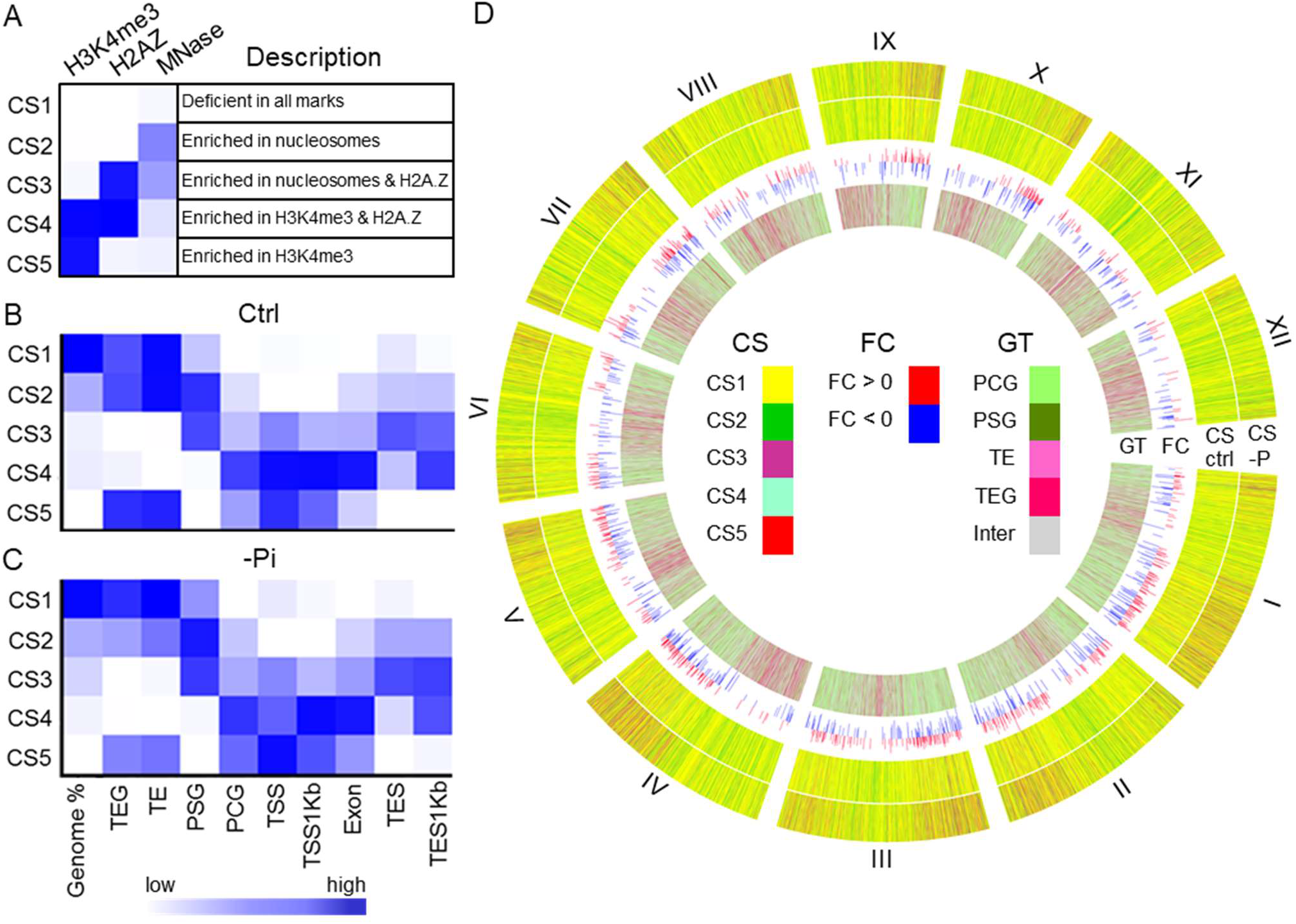
Chromatin state predictions for control (Ctrl) and Pi deficiency (–Pi) samples defined by H3K4me3, H2A.Z and nucleosome occupancy. (A) Emission parameters for 5 chromatin states (CS). The darker blue color corresponds to a greater probability of observing the mark in the state. Overlap fold enrichment of various genomic regions with 5 chromatin states in Ctrl (B) and –Pi (C) samples. PCG, protein coding genes; PG, pseudogenes; TE, non-expressed transposable element-related genes; TEG, expressed transposable element-related genes; TSS, transcription start site; TES, transcription termination site; TSS1Kb, 200 bp upstream to 800 bp downstream of the TSS; TES1Kb, 800 bp upstream to 200 bp downstream of the TES. (D) Circos plot showing the chromatin states (in 5kbp bins) of the whole genome. The first and second rings show the chromatin state in -Pi and Ctrl condition respectively. The third ring shows fold change of differentially expressed genes and the last ring represents four gene types. Each segment in circus plot showed one chromosome (Chr) in the rice genome.

### Pi starvation has a dramatic impact on chromatin structure

To characterize the impact of Pi starvation on chromatin structure we compared the distribution of chromatin states between control and Pi-deficiency (–Pi) conditions. First we measured the genome-wide coverage changes for each chromatin state by calculating the fold change in the total number of genomic bins in the –Pi sample relative to the control. As shown in Supplemental Figure S3, the –Pi sample had 2.1-fold more CS3 and 1.4-fold less CS4 compared to control. This suggested a global increase of H2A.Z and/or decrease of H3K4me3. Next we analyzed the enrichment of each chromatin state within the four gene types (Figure 3C,D). In response to Pi starvation, TE and TEG increased in CS1, but decreased in CS2 and CS5, consistent with a loss of H3K4me3. On the other hand, PSG and PCG did not exhibit any major shifts overall in response to Pi deficiency, but at the TSS of PCG, CS4 decreased and CS5 increased. Also, at the TES proximal region of PCG, CS4 decreased and CS3 increased. Together this reveals an overall trend whereby, at PCG, Pi deficiency leads to decreased H2A.Z at the TSS and decreased H3K4me3 near the TES.

To examine the chromatin state transitions of PCG in more detail, we compared the chromatin state of each PCG at its TSS (i.e. the 200-bp bin containing the TSS) in control and –Pi samples (Figure 4). Over 40 % of PCG exhibited a transition at their TSS in response to Pi starvation. The largest groups of transitions were CS4 to CS3 (n = 4,088), CS4 to CS5 (n = 2,355), and CS5 to CS1 (n = 2,496). Gene Ontology (GO) enrichment analysis showed significantly enriched GO terms (FDR < 0.05) for eight of the transition groups (Supplemental Dataset 2). Because of the inherent redundancy of GO term enrichment analysis, we used GOMCL (Wang et al., in review) to enhance the functional annotations of the enriched GO terms for the three largest groups of CS transitions. GOMCL employs Markov Clustering to identify clusters of GO terms based on the proportion of overlap among terms. As shown in Figure 4, the enriched GO terms for CS4-CS3 genes fell into five GOMCL clusters, including transcription factor activity, response to endogenous stimulus, cell wall, oxygen binding, and response to extracellular stimulus. In contrast, CS4-CS5 genes were enriched in GO terms defined by nine GOMCL clusters, which among other functional categories, were related to translation and gene expression, nuclear functions, plastid functions, nucleic acid metabolism, development, and RNA binding. Interestingly, CS5-CS1 genes shared essentially the same enriched GO terms (Supplemental Dataset 2) and GOMCL clusters (Figure 4) as CS4-CS5 genes. One explanation for this is that the CS5-CS1 and CS4-CS5 transitions are frequently found together at the TSS. Indeed, examination of the bins that flank the TSS (Supplemental Figure S4) showed that CS5-CS1 genes were approximately four times more likely than random to exhibit a CS4-CS5 transition in the bin downstream of the TSS (p-value < 0.001). Similarly, the CS4-CS5 genes were 3.5-fold more likely to contain a CS5-CS1 transition upstream of the TSS (p-value < 0.001). In contrast, CS5-CS1 genes with a CS4-CS5 upstream bin, and CS4-CS5 genes with a CS5-CS1 downstream bin were similar to random or under-represented, respectively. Thus, the identification of subgroups of functionally similar genes with CS5-CS1 and CS4-CS5 transitions at their TSS is reflective of these genes containing a specific pair of transitions (CS5-CS1 + CS4-CS5, 5’-3’) at the TSS.

**Figure 4.**
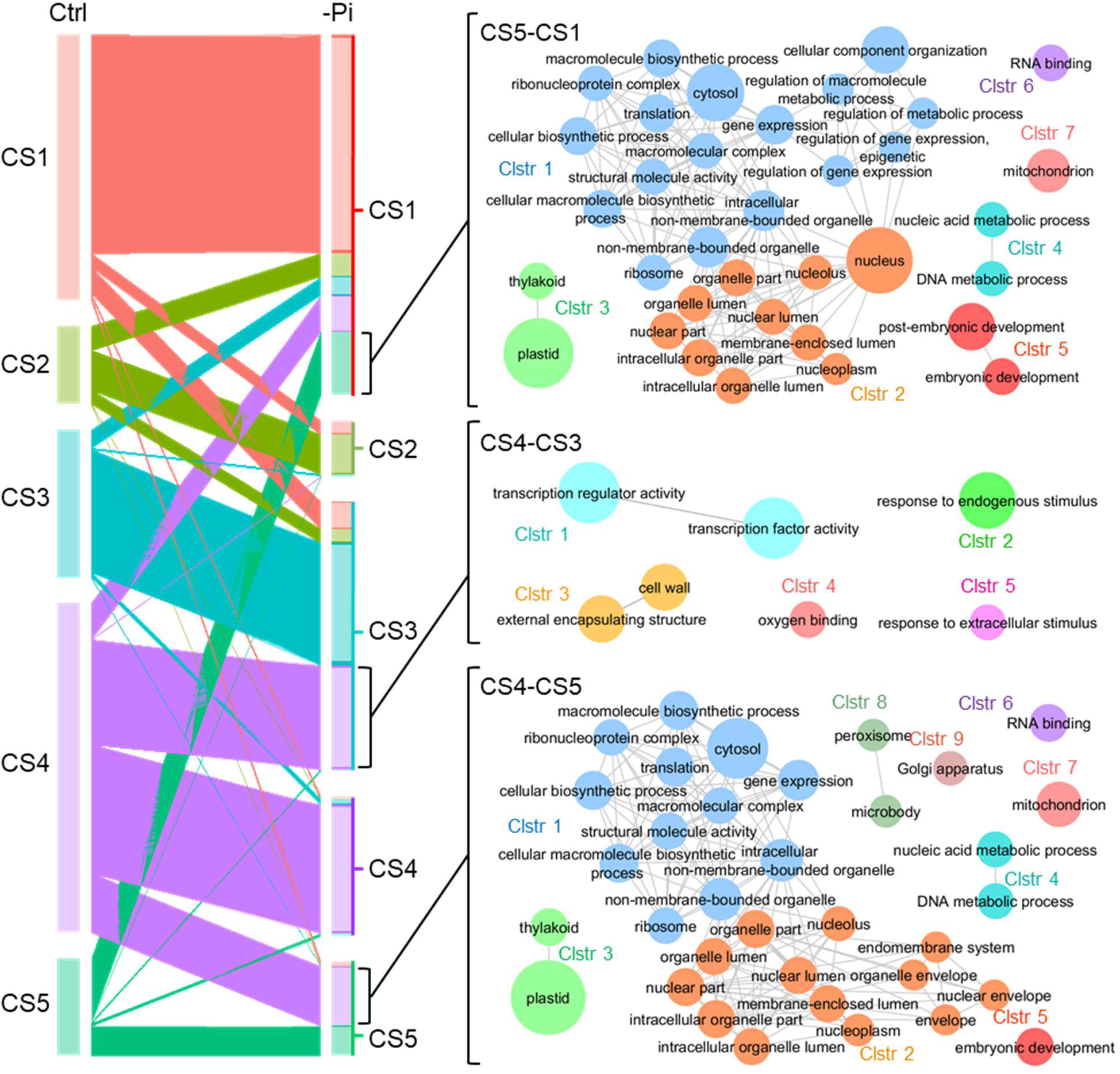
Chromatin state (CS) transitions of protein-coding genes from control (Ctrl) to Pi deficiency (-Pi) conditions. (Left) The size of the segment represents the number of gene in each CS and the width of the ribbons represent the number of genes with a transition to another CS. (Right) Networks representing Gene Ontology Markov Clustering (GOMCL) terms enriched in CS5-CS1, CS4-CS3 and CS4-CS5 groups. Cytoscape was used to visualize enriched GO terms.

### Chromatin state transitions correlate with differential expression of Pi deficiency-responsive genes

We analyzed our recent RNA-seq experiments (Zahraeifard et al., 2018; Supplemental Table S1) to investigate the relationship between gene expression and chromatin state transitions in response to Pi starvation. Differential expression analysis with DESeq2 identified 1385 differentially-expressed genes (DEGs) in response to Pi starvation, 694 up-regulated and 691 down-regulated (adjusted P-value < 0.001; Supplemental Figure S5, Supplemental Dataset 3) GO terms enriched for up-regulated genes included response to stress, lipid metabolic process, and signal transduction, whereas down-regulated genes were enriched in growth, cell-cell signaling, and lipid, carbohydrate, and secondary metabolic processes (Supplemental Table S2, Supplemental Figure S5B,C). Although lipid metabolism was overrepresented in both groups of DEGs, genes linked to carotenoid biosynthesis and alpha-Linolenic acid metabolism were among the up-regulated DEGs, whereas cutin, suberin, and wax biosynthesis were among the down-regulated DEGs. Overall, the functional categories of these DEGs were similar to those from previous transcriptome studies of Pi-deficient plants (Thibaud et al., 2010, Cai et al., 2013, Secco et al., 2013, Zahraeifard et al., 2018).

To determine whether any chromatin state transitions were over- or under-represented among the DEGs, we quantified the significance of overlap via bootstrapping analyses (1000 iterations; binomial test, p-value < 0.001; Figure 5). These analyses revealed several significant biases between CS transitions and DEGs. First, down-regulation of gene expression correlated with a gain of H2A.Z, as indicated by enrichment of down-regulated genes with CS1-CS3 and CS2-CS3 transitions and under-representation of up-regulated genes with a CS2-CS3 transition. Reciprocally, up-regulation of gene expression correlated with a loss of H2A.Z, as indicated by enrichment of up-regulated genes among CS3-CS1 genes. These observations support a role for H2A.Z as a repressive chromatin mark during Pi starvation, in which some genes are repressed by the deposition of H2A.Z, whereas other genes are induced (de-repressed) by its removal. Second, genes containing H2A.Z and H3K4me3 that exhibited decreases in both marks in response to Pi deficiency (CS4-CS1) were also enriched among up-regulated genes. This suggests a negative role for not only H2A.Z, but also H3K4me3, in which the loss of both marks from this group of genes results in their de-repression. Third, up- and down-regulated DEGs were both enriched among CS4-CS3 transition genes (i.e. those with a decrease in H3K4me3 but maintenance of H2A.Z). Interestingly, this suggests a possible dual role of H3K4me3 in Pi-responsive gene modulation. Finally, the other two prominent groups of transitions, CS5-CS1 and CS4-CS5, which contain many translation-related genes, were under-represented among down-regulated DEGs. This indicates that genes exhibiting these transitions, or pair of transitions (Supplemental Figure S4), at the TSS are unlikely to be differentially expressed after 24-hours of Pi deficiency. Because a number of translation-related genes were previously shown to be down-regulated by long-term (21-day) Pi deficiency in rice shoots (Secco et al., 2013), we carried out a bootstrapping analysis to test whether our CS5-CS1 and CS4-CS5 genes were enriched among those DEGs. Indeed, both CS5-CS1 and CS4-CS5 groups were enriched (p-value < 0.01) among genes down-regulated by long-term Pi deficiency (Supplemental Figure S6). This suggests that the chromatin dynamics observed at these genes after 24 hours of Pi starvation is a prelude to decreased transcript abundance not observable until after a longer duration of Pi deficiency.

**Figure 5.**
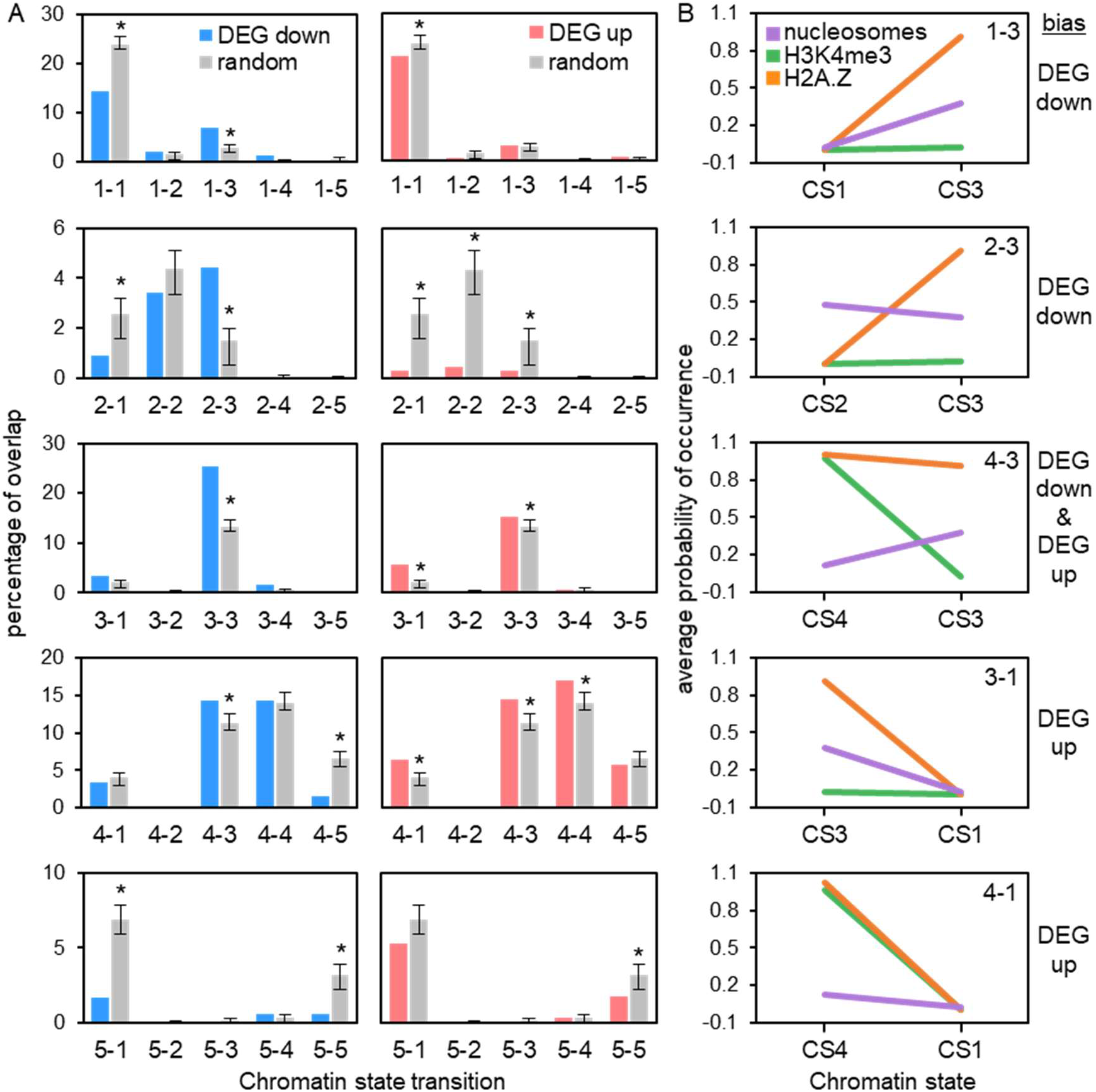
Chromatin state (CS) transitions are associated with differentially-expressed genes (DEGs) under phosphate deficiency. (A) Bootstrapping analysis showing the overlap between genes exhibiting chromatin state transitions and down-regulated or up-regulated genes in response to Pi deficiency. Data are means (±SD) for 1000 iterations. (B) Values are the average probability of each chromatin mark at the CS shown. The category of DEG (up or down) that is biased to the CS is shown at right.

In addition to the biases between DEGs and chromatin state transitions, there were also biases to groups of genes that did not transition (Figure 5). Both up- and down-regulated DEGs were significantly enriched among CS3 genes that did not transition (i.e. CS3-CS3), and were under-represented among CS1-CS1 and CS5-CS5 genes. Furthermore, up-regulated DEGs were enriched among CS4-CS4 genes and under-represented among CS2-CS2 genes. These results show that responsive genes are likely to contain H2A.Z, which is consistent with previous reports (Coleman-Derr and Zilberman, 2012, Zahraeifard et al., 2018). Taken together, these biases demonstrate that specific chromatin dynamics at the TSS are linked to subsets of genes differentially expressed by Pi starvation.

### Differentially-expressed genes exhibiting a CS4 to CS3 chromatin transition suggest a coordinated Pi-deficiency regulatory network targeting the apoplast

As shown above, the largest group of genes exhibiting a chromatin state shift in response to Pi deficiency was the CS4-CS3 group (Figure 4, Supplemental Dataset 2). These genes were also significantly enriched among both up- and down-regulated DEGs (Figure 5), and GO term enrichment analysis of the DEGs indicated functions linked to the cell wall, responses to biotic stress, and catalytic activity (Supplemental Figure S7). Due to the relatively limited GO term assignments for rice loci, we carried out extensive data mining on the CS4-CS3 DEGs, which allowed us to assign putative functional and subcellular localization information to 178 (91%) of the 196 DEGs (Supplemental Dataset 4). These DEGs encode components with putative functions in signal transduction (37%), cell wall structure (23%), lipid composition (13%), transcription regulation (10%), secondary metabolism (9%), primary metabolism, or cell growth (3%), which are mostly targeted to the apoplast (31%), plasma membrane (28%), nucleus (18%), cytosol (10%), or plastid (6%) (Figure 6A). Strikingly, more than half (53%) of the CS4-CS3 DEGs are predicted to encode proteins targeted to the apoplast or plasma membrane, and have functions in signaling or cell wall and lipid composition. Among this group are a number of pectinases, arabinogalactan proteins (AGPs), and expansins that mostly are down-regulated by the 24-hour Pi deficiency treatment (Figure 6B, Supplemental Dataset 4). A previous study in Arabidopsis identified a similar response of cell wall hydrolytic enzyme-encoding loci in roots subjected to Pi-deficiency treatments of 1, 6, and/or 24 hours (Lin et al., 2011). Together this suggests that modification of the cell wall is an early and prominent response to Pi starvation in roots and shoots across plant species. In addition to the down-regulation of cell wall-related components was a large group of signaling components, including many receptor-like kinases (RLKs), that were predominantly up-regulated (Figure 6B, Supplemental Dataset 4). One of the RLKs is a *Catharanthus roseus* RLK1-like kinase orthologous to Arabidopsis FERONIA (FER), which has been shown to regulate cell expansion in response to diverse developmental and environmental cues (Liao et al., 2017). For example, during salinity stress, FER maintains cell wall integrity, and is necessary for root growth recovery (Feng et al., 2018). Recently it was demonstrated that FER is one component of a signaling module that transduces cell-wall signals during salt stress (Zhao et al., 2018). In the absence of salt stress, a group of apoplastic leucine-rich repeat extensins (LRX) bind to RAPID ALKALINIZATION FACTOR (RALF) peptides. In response to salt stress, LRX and RALF dissociate, and RALF peptides bind FER. This results in FER internalization and, subsequently, inhibition of growth and initiation of stress responses. Calcium transients and SITE-1 PROTEASE (S1P) activity also play roles in RALF/FER signaling (Stegmann et al., 2017, Feng et al., 2018). Notably, our CS4-CS3 DEG list also contains genes encoding six RALF peptides (out of 14; (Campbell and Turner, 2017) an LRX, several Ca2+ transport-related components (e.g. Ca2+ ATPase and calmodulin), and two S1P proteases (Supplemental Dataset 4). In addition to the signaling and cell wall components were a number of transcription factors among the CS4-CS3 DEGs, including five AP2 superfamily factors, two HLH factors, and two WRKY transcription factors. These represent families of transcription factors known to be responsive to a number of biotic and abiotic stressors. It is tempting to speculate that these regulatory genes, along with the differentially-expressed CS4-CS3 structural genes comprise a transcriptional regulatory network aimed at transducing Pi deficiency signals and initiating reduced cellular growth and tolerance to low Pi (Figure 7).

**Figure 6.**
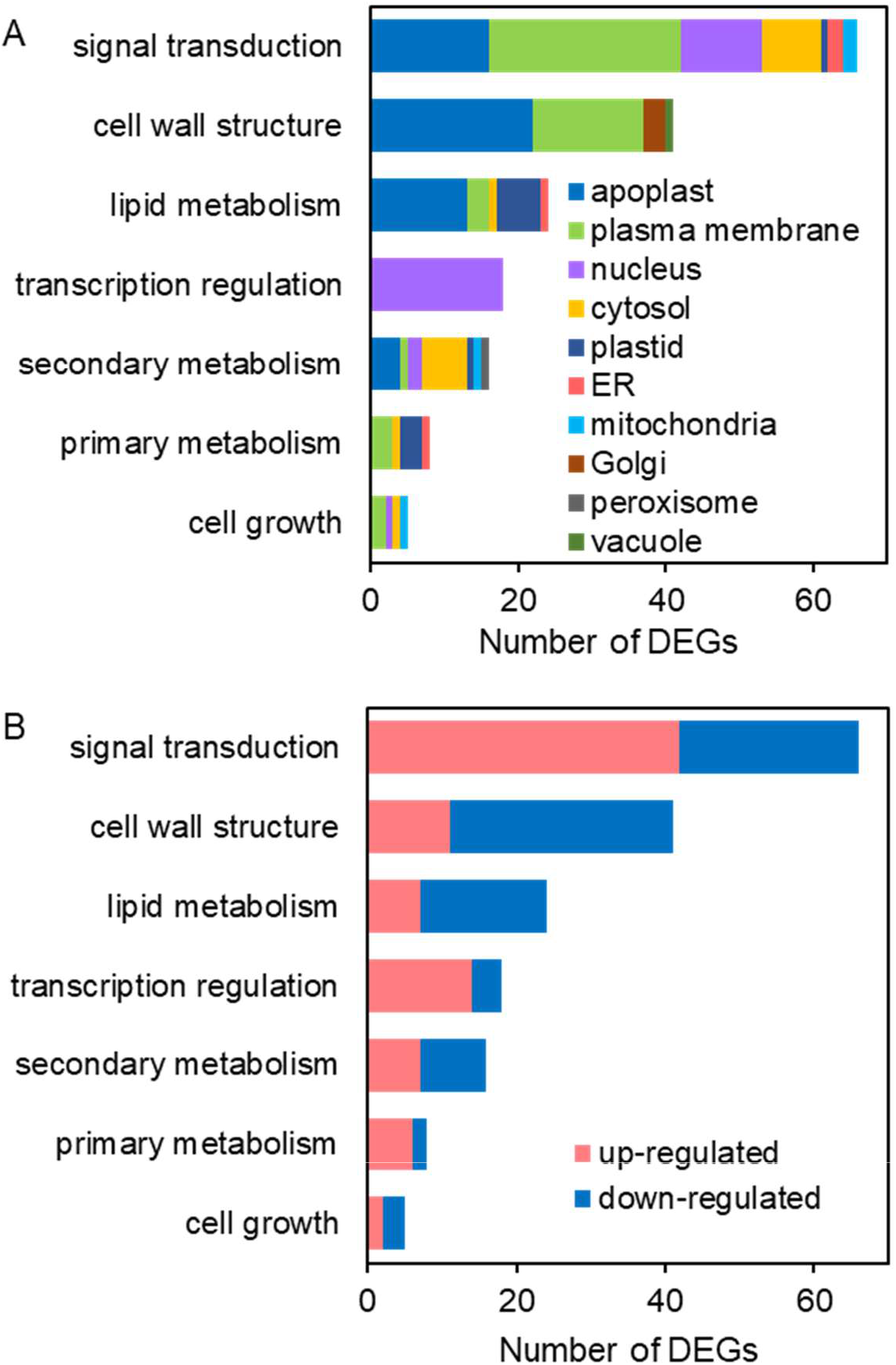
Predicted functions and subcellular locations of differentially-expressed genes (DEGs) having a chromatin state (CS) transition of CS4 to CS3.

**Figure 7.**
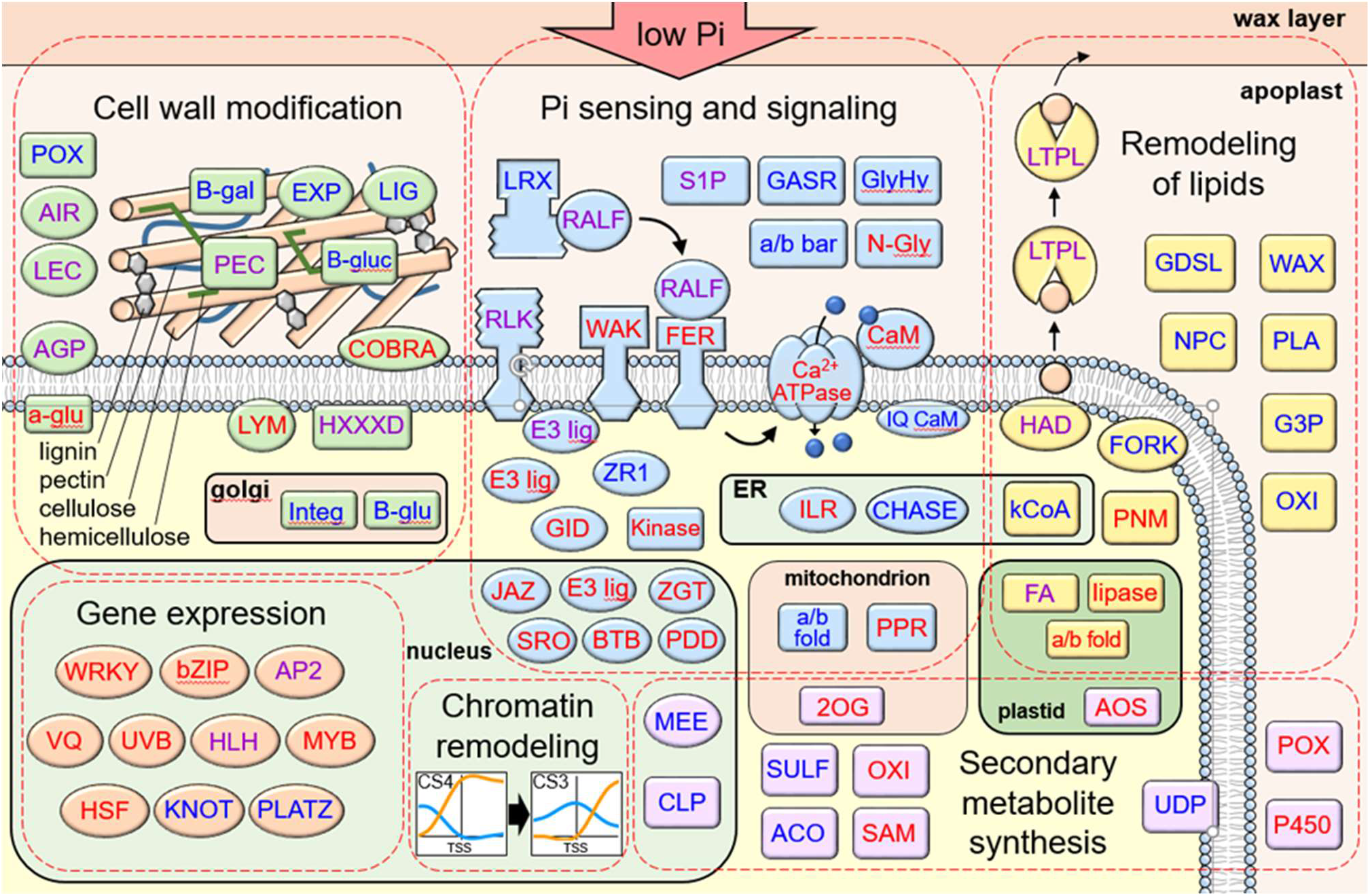
Predicted interactions and functions of differentially-expressed genes having a chromatin state (CS) transition of CS4 to CS3. Abbreviations: 2OG, 2OG-Fe oxygenase; a/b bar, A/B barrel; a/b fold, alpha/beta fold hydrolase; ACO, 1-aminocyclopropane-1-carboxylate oxidase; a-glu, heparan-alpha-glucosaminide N-acetyltransferase; AGP, arabinogalactan protein; AIR, auxin response protein; AOS, allene oxide synthase; AP2; B-gal, beta-galactosidase; B-glu, Beta glucan synthase; B-gluc, beta-glucuronidase; BTB, Bric-a-Brac, Tramtrack, Broad Complex protein; bZIP; Ca ATPase; CaM, Calmodulin-related calcium sensor; CHASE; CLP, ATP-dependent caseinolytic protease/crotonase; COBRA, AtCOBRA-like; E3 lig, ubiquitin ligase; EXP, expansin; FA, fatty acid hydroxylase; FER, AtFERONIA ortholog; FORK, FORKED1-like; G3P, glycerol-3-phosphate acyltransferase; GASR, GASA/GAST/Snakin; GDSL, GDSL-like lipase/acylhydrolase; GID, gibberellin receptor; GlyHy, glycosyl hydrolase; HAD, HAD phosphoethanolamine/phosphocholine phosphatase; HLH; HLH helix-loop-helix transcription factor; HSF, heat shock factor; HXXXD, HXXXD-type acyl-transferase; ILR, IAA-amino acid hydrolase; Integ, cell wall integrity protein; IQ CaM, IQ calmodulin-binding motif protein; JAZ, ZIM domain-containing JAZ protein; kCoA, 3-ketoacyl-CoA synthase; kinase; KNOT, knotted-1-like homeobox protein; LIG, lignin dirigent; lipase; LRX, leucine-rich repeat extensin; LTPL, Protease inhibitor/seed storage/LTP protein; LYM, lysM domain-containing GPI-anchored protein; MEE, maternal effect embryo arrest; MYB; N-Gly, shiga/ricin-like N-glycosidase; NPC, non-specific phospholipase; OXI, oxidoreductase; P450, cytochrome P450; PDD, PD-(D/E)XK nuclease superfamily protein; PEC, pectinase; PLA, phospholipase A; PLATZ; PNM, phosphoethanolamine N-methyltransferase; POX, peroxidase; PPR, pentatricopeptide repeat protein; RALF, Rapid ALkalinization Factor; RLK, receptor-like kinase; S1P, Subtilisin Site-1 Protease; SAM, S-adenosyl-L-methionine-dependent methyltransferases; SRO, OsSRO1c; SULF, sulfotransferase; UDP, UDP-glucuronosyl/UDP-glucosyltransferase; UVB, ultraviolet-B-repressible protein; VQ, VQ domain containing protein; WAK, wall-associated kinase; WAX, WAX2-like; WRKY; ZGT, ZGT circadian clock coupling factor; ZR1, FYVE zinc finger domain protein.

## Discussion

### H3K4me3 and H2A.Z exhibit overlapping and divergent localization patterns

Despite being widely recognized as marks of active transcription, assigning specific roles for H3K4me3 and H2A.Z in regulating transcription has been challenging. For instance, H3K4me3 is often assumed to promote transcription, but loss or severe depletion of H3K4me3 levels results in relatively few gene expression changes (Clouaire et al., 2012, Margaritis et al., 2012). Also, whereas loss of H3K4me3 at most genes has no impact on expression, H3K4me3 has been linked to both activation and repression of subsets of genes (Weiner et al., 2015, Cano-Rodriguez et al., 2016). Like H3K4me3, H2A.Z is often associated with gene activity, but plays a complex role in modulating gene expression. Evidence indicates that H2A.Z acts to both promote and repress gene expression, depending on the environmental or developmental context, genic location, and relevant loci (Deal et al., 2007, Zilberman et al., 2008, March-Díaz and Reyes, 2009, Kumar and Wigge, 2010, Smith et al., 2010, Berriri et al., 2016, Sura et al., 2017, Zahraeifard et al., 2018). Interactions among multiple chromatin modifications add complexity to identifying specific chromatin-level mechanisms that modulate gene expression, particularly in light of contradictory findings. For example, Arabidopsis H2A.Z has been proposed to facilitate H3K4 trimethylation at miR156 loci (Xu et al., 2018) but antagonize H3K4me3 abundance at anthocyanin biosynthetic genes (Cai et al., 2019). Thus there is a need to investigate multiple aspects of chromatin structure in order to gain insight into chromatin-level mechanisms that impact gene expression.

Herein we used ChromHMM to combine our H3K4me3 ChIP-Seq data from this study with our previous H2A.Z ChIP-Seq (Zahraeifard et al., 2018) and MNase-Seq (Zhang et al., 2018) data to define 5 chromatin states (CS1-CS5) in rice shoots. Genic regions were enriched in CS4, which is characterized by moderate nucleosome occupancy and relatively high levels of H2A.Z and H3K4me3. The TSS of protein-coding genes were also enriched in CS4, as well as CS3 and CS5, which contain only H2A.Z or H3K4me3, respectively. In contrast, the TES of protein-coding genes was only enriched in CS3. This suggests that H3K4me3 functions mostly at the TSS, whereas H2A.Z functions across the gene. This is generally consistent with previous reports on the functions of H3K4me3 and H2A.Z. Studies in a number of organisms have shown that H3K4me3 localizes near the TSS of active protein-coding genes (Santos-Rosa et al., 2002, Liu et al., 2005, Bernstein et al., 2006, Barski et al., 2007, Zhang et al., 2009, Van Dijk et al., 2010, Du et al., 2013, Zong et al., 2013). Our data further support this by showing a prominent peak of H3K4me3 at the TSS of rice PCG (Figure 1A) that is positively correlated with basal gene expression (Figure 2). H2A.Z is also abundant at the TSS of PCG, but appears to play roles in gene expression by localizing to gene bodies and the TES, as well (Coleman-Derr and Zilberman, 2012, Sura et al., 2017, Zahraeifard et al., 2018). In contrast to H3K4me3, TSS-localized H2A.Z is negatively correlated with basal expression (Figure 2; (Zilberman et al., 2008, Coleman-Derr and Zilberman, 2012, Yelagandula et al., 2014, Dai et al., 2017, Zhang et al., 2017, Zahraeifard et al., 2018). Interestingly, our data show that both H2A.Z and H3K4me3 localized downstream of the TSS region are negatively correlated with expression. Previous studies have reported this phenomenon for H2A.Z (Zilberman et al., 2008, Coleman-Derr and Zilberman, 2012, Sura et al., 2017), but Arabidopsis H3K4me3 was shown to be positively regulated with expression (Van Dijk et al., 2010). This may reflect a difference in the role of H3K4me3 at the 3’ genic region in different plant species. On the other hand, a H3K4me3 profile of genes from an allotetraploid cotton genotype generally showed a negative correlation with expression, whereas a diploid cotton genotype in the same study exhibited a positive correlation (You et al., 2017). H3K4me3 at the TES was reported to play a role in modulating antisense transcription, thereby repressing sense transcription (Ponting et al., 2009, Cui et al., 2012). Therefore, genotypic or cell type-dependent differences in antisense transcription may contribute to the correlation of TES-localized H3K4me3 with sense transcription. Further investigation is required to understand the nature of the differences in 3’ H3K4me3-dependent regulation of gene expression across samples and species.

### Pi-starvation induced chromatin dynamics correlate with gene repression and induction

Often, the disruption of chromatin remodelers, such as H3K4 methyltransferases, through mutagenesis do not have substantial impacts on global steady-state transcription (Guo et al., 2010, Chen et al., 2017, Howe et al., 2017). On the other hand, a number of studies have identified significant roles for particular chromatin remodelers in differential expression in response to environmental stimuli (Ding et al., 2011, Ding et al., 2012, Weiner et al., 2015). Our data support this by revealing that more than 40% of all rice PCG in shoots exhibit a chromatin state transition at their TSS in response to a 24-hour Pi deficiency treatment, and that several specific transitions strongly correlate with subgroups of genes differentially-expressed by Pi starvation. Indeed, our results suggest that multiple chromatin remodelers are responsive to Pi deficiency and influence expression of distinct subsets of target genes.

Genes with CS1-CS3 or CS2-CS3 transitions exhibit increases in nucleosome occupancy and H2A.Z deposition in response to Pi starvation, and are enriched in down-regulated genes, whereas CS3-CS1 genes, which exhibit decreases in nucleosome occupancy and H2A.Z, are enriched in up-regulated genes. These correlations are consistent with a repressive role for H2A.Z at the TSS in modulating Pi deficiency response genes. This is consistent with our recent work in rice (Zahraeifard et al., 2018) and previous reports in Arabidopsis (Dai et al., 2017, Sura et al., 2017), which all provide evidence for H2A.Z acting as a repressor of expression when localized at gene bodies or the TSS. Work in Arabidopsis also showed general co-localization of H2A.Z and H3K4me3 in promoter regions, but a negative correlation of the two marks at the TSS of genes exhibiting relatively high H2A.Z, as well as a positive correlation between nucleosome occupancy and H2A.Z at the +1 nucleosome, suggesting that H2A.Z deposition at the +1 nucleosome is linked to high nucleosome occupancy, low H3K4me3, and low gene accessibility (Dai et al., 2017). Our data bolster support for a model where H2A.Z at the TSS, likely the +1 nucleosome, regulates a subset of Pi-deficiency response genes that contain low H3K4me3 and relatively low basal expression. In response to Pi starvation, H2A.Z is either removed or deposited, resulting in de-repression (CS3-CS1) or repression (CS1-CS3/CS2-CS3), respectively. Similar to CS3-CS1, genes with a CS4-CS1 transition, which exhibit a loss of both H2A.Z and H3K4me3, are enriched in up-regulated genes (Figure 6). These genes tend to be more highly expressed during control conditions than CS3 genes, and therefore have a stronger requirement for H3K4me3 for basal expression. In response to Pi starvation, the combined loss of H2A.Z and H3K4me3 may reflect some dependence of H3K4me3 on H2A.Z at these genes, similar to how H2A.Z was suggested to facilitate H3K4me3 deposition at two miR156-encoding genes in Arabidopsis (Xu et al., 2018).

Among the gene groups that exhibit chromatin state transitions, the CS4-CS3 group contains the largest number of genes, and is characterized by a loss of H3K4me3, but maintenance of H2A.Z, during Pi starvation. Interestingly, these genes are enriched among both up- and down-regulated genes, indicating that loss of H3K4me3 is linked to gene activation and repression during Pi deficiency. In contrast to H2A.Z, H3K4me3 is generally not recognized as playing a negative role in gene expression. Studies in a variety of plant species and tissues have examined the change in genic levels of H3K4me3 in response to environmental stressors (Tsuji et al., 2006, Sokol et al., 2007, Kim et al., 2008, Van Dijk et al., 2010, Jaskiewicz et al., 2011, Zeng et al., 2019). These studies generally reported increases in H3K4me3 at genes up-regulated by stress. However, most of the studies examined relatively small numbers of genes, and the genome-level studies that compared average H3K4me3 genic profiles between control and stressed samples actually found substantial decreases in 5’ localization of H3K4me3 in response to stress (Zong et al., 2013, Zeng et al., 2019). We observed a similar effect when comparing the H3K4me3 profiles for all PCG between control and Pi deficiency conditions (Figure S2). One explanation for our CS4-CS3 genes being linked to both induction and repression is that the TSS of the corresponding genes contain bivalent histone modifications. Bivalent domains are characterized by containing both active and repressive histone modifications. First described in mouse embryonic stem cells were bivalent domains containing H3K4me3 and H3K27me3, in which H3K4me3 is proposed to poise genes for activation, whereas H3K27me3 maintains the genes in a repressed state (Bernstein et al., 2006). A recent study in potato tuber found an association between genes containing the bivalent H3K4me3 and H3K27me3 marks and differential expression in response to cold stress (Zeng et al., 2019). Interestingly, the bivalent mark was enriched among up-regulated genes linked to stress responses, as well as down-regulated genes linked to developmental processes. The authors proposed that the bivalent H3K4me3-H3K27me3 domain confers greater accessibility to regulatory proteins that can induce or repress genes in response to cold stress. A similar phenomenon might explain our observed link between CS4-CS3 genes and both up- and down-regulation of genes in response to Pi starvation. A decrease in H3K4me3 at the TSS may reflect a switch from nucleosomes modified with only H3K4me3 to nucleosomes containing both H3K4me3 and H3K27me3. This would favor enhanced DNA accessibility, which could facilitate the targeting of transcriptional machinery for induction or repression by the appropriate transcriptional machinery. Recently, an interaction between H2A.Z deposition and H3K27 tri-methylation was reported in Arabidopsis, in which H2A.Z deposition promotes H3K27 tri-methylation (Carter et al., 2018). It is possible that the maintenance of H2A.Z at the CS4-CS3 genes is required for proper H3K27me3 deposition at the bivalent marks. Future experiments that examine H3K27me3 localization would shed light on this hypothesis.

### Differential expression of cell wall-related genes correlates with decreased H3K4me3 and maintenance of H2A.Z

Cell walls provide rigidity to plant cells but are also restrictive to cell expansion. Thus, cells must simultaneously weaken cell wall structure and maintain turgor and cell integrity to achieve growth (Voxeur et al., 2016). Correspondingly, plants must employ signaling mechanisms aimed at regulating cell wall structure in response to developmental and environmental cues. Several plasma-membrane localized receptor-like kinases, such as FERONIA (FER), have been implicated in cell-wall integrity sensing in response to a variety of environmental stressors (Liao et al.,. 2017). The majority of our CS4-CS3 DEGs encode putative apoplastic or plasma membrane proteins with predicted roles in signaling and cell wall composition. The signaling genes were mostly up-regulated, whereas the cell wall related genes were largely down-regulated. Comparing the transcriptomic profile of the CS4-CS3 DEGs to public transcriptome studies using Genevestigator (Hruz et al., 2008) revealed substantial overlap with several pairwise comparisons from a previous study on rice lamina joint development (Zhou et al., 2017). Comparisons between older stages of development (maturation or post-maturation) with a younger stage showed similar expression profiles as our CS4-CS3 Pi-deficiency DEGs (not shown). Interestingly, cell-wall thickening is a prominent feature during younger stages of lamina joint development, and this declines over time. This may suggest that Pi starvation results in decreased cell wall thickening, or more generally, a decrease in cell elongation. Transcriptomic profiles of several biotic and abiotic (e.g. salinity and heat) stressors also showed high similarity to our CS4-CS3 DEG profile, suggesting the apparent apoplastic signaling network overlaps with multiple stressors. Our CS4-CS3 DEG list contains many orthologs of Arabidopsis components involved in salinity stress responses, including FER, LRX, RALF (Zhao et al., 2018). It is of interest to evaluate whether the rice orthologs exhibit similar functions in response to stressors including salinity and Pi starvation.

### A distinct pair of chromatin state transitions may poise translation-related genes for repression

Following the CS4-CS3 gene group, the transitions with the most genes were the CS5-CS1 and CS4-CS5 transitions, which were enriched with similar functional categories of genes including those related to translation, particularly a number of ribosomal protein genes (Figure 4). Examination of the two bins adjacent to the TSS revealed that a number of these genes contained both transitions with the CS5-CS1 transition immediately upstream of the CS4-CS5 transition. Our bootstrapping results showed that these genes are not enriched among our DEGs. On the contrary, the CS4-CS5 subgroup are under-represented among down-regulated DEGs (Figure 6). Interestingly, a group of genes shown in a previous study (Secco et al., 2013) to be down-regulated after 21 days of Pi deficiency were enriched among our CS5-CS1/CS4-CS5 genes (Supplemental Figure S6). This might indicate that 24 hours of Pi deficiency is sufficient to observe chromatin dynamics at these genes without observing a corresponding, detectable decline in transcript abundance. We propose that the sequential CS5-CS1 and CS4-CS5 transitions observed at the TSS reflect genes under control conditions that contain low H2A.Z and high H3K4me3 in the -1 nucleosome and high levels of both marks in the +1 nucleosome. Pi starvation, then, results in a moderate loss of nucleosome occupancy at both the +1 and -1 nucleosomes, and specific removal of H3K4me3 from the -1 nucleosome and H2A.Z from the +1 nucleosome. In yeast, Spp1 promotes the H3K4 trimethylase activity of the Set1 complex (Morillon et al., 2005). As a result, deletion of Spp1 results in substantial loss of global H3K4me3 levels, but the remaining H3K4me3 (approximately 20 %) is not evenly distributed among genes. Genes that retain the highest levels of H3K4me3 in *Δspp1* mutants are enriched in ribosomal protein genes and other translation-related genes, whereas genes exhibiting the most severe H3K4me3 depletion are enriched in stress-related genes (Howe et al., 2014). Also, the Spp1-independent genes tend to be more highly expressed during control conditions, and repressed during environmental stress, whereas the Spp1-dependent genes generally exhibit low expression during control conditions and induced expression during stress. Finally, in response to diamide stress, many yeast ribosomal protein genes are down-regulated and exhibit a decrease in H3K4me3 (Weiner et al., 2015). Our data suggest that rice employs different mechanisms to modulate H3K4me3 levels at distinct gene groups, similar to yeast. This is consistent with our CS4-CS3 and CS5-CS1 gene groups undergoing decreases in H3K4me3 via different chromatin remodeling complexes. Future studies on the roles of H3K4me3 and H2A.Z, in conjunction with additional marks such as H3K27me3, in the Pi deficiency-dependent regulation of gene expression will provide valuable information on the chromatin dynamics that impact low-Pi adaptation mechanisms.

## Materials and methods

### Plant material and growth conditions

Sterilization and pre-germination (1 day at 37 °C followed by 2 days at 28 °C) were carried out on rice cultivar Nipponbare (*Oryza sativa ssp. japonica*) seeds. Seeds were transferred to 12-h light/12-h dark, at 30 °C/22 °C condition to germinate for 14 days. Seedlings were grown hydroponically in modified Yoshida Rice culture media as described (Yoshida et al., 1971, Secco et al., 2013). The solution was replaced every 7 days. After 21 days, seedlings were used for a 24-hour Pi-deficiency treatment (modified Yoshida Rice solution without NaH_2_PO_4_).

### ChIP-seq

Four grams of frozen shoot tissue from 24-hour Pi deficiency or control treatment was used to perform chromatin immunoprecipitation (ChIP) as previously described (Zahraeifard et al., 2018) using the antibody (Millipore; lot number 2648189) against H3K4me3 and input genomic DNA as a control. Three biological replicates were used for both input and antibody treatments. Purification of ChIP DNA was carried out with the Clean and Concentrator kit (Zymo Research). Libraries were constructed using 1:20 diluted adaptor from Kapa Biosystems Hyper Library Construction Kit and 10 cycles of DNA amplification. Libraries were quantitated (qPCR) and multiplexed, and single-end sequencing was completed with a HiSeq2500 (Illumina) using a HiSeq SBS sequencing kit (version 4) for 101 cycles at the University of Illinois Roy J. Carver Biotechnology Center. Approximately 147 million ChIP-seq reads were quality-checked and cleaned using FastQC and Trimmomatic-0.33 (Andrews, 2010, Bolger et al., 2014). Using Bowtie, the reads were aligned to MSU Rice Genome Annotation Release 7.1 (MSU7.1) with one mismatch allowed to retain uniquely mapped reads. The SICER software package (Zang et al., 2009) was used to define the H3K4me3 enrichment regions with the following parameters (W = 200, G = 200, FDR < 1.00E-02). The input genomic DNA was used as a background control. Differential H3K4me3 enrichment peaks between control and Pi deficiency samples were determined using SICER-df.sh shell script (W = 200, G = 200, FDR < 1.00E-02). We defined the existence of peaks with protein-coding genes (PCG) if 50 % of peaks overlapped with PCG (including 250 bp upstream and downstream) using BEDTools intersect (Quinlan and Hall, 2010). The genome-wide distribution pattern of H3K4me3 and the published profile of H2A.Z (Zahraeifard et al., 2018) were visualized using ngs.plot (Shen et al., 2014). K-means clustering within ngs.plot was used to find different patterns of H3K4me3. Gene ontology (GO) terms enriched among clusters were analyzed with BiNGO and visualized with Cytoscape (Maere, Heymans et al. 2005).

### RNA-seq analysis

RNA-sequencing reads were generated previously (Zahraeifard et al.., 2018). A minimum of 58 million high-quality RNA-seq reads (100-bp single end) per sample were mapped to the MSU Rice Genome Annotation Release 7.1 (MSU7.1) using Bowtie2 tools (Langmead and Salzberg, 2012). Fragments per kilobase of transcript per million mapped reads (FPKM) were calculated with the Cuffdiff tool (Trapnell et al., 2012). DESeq2 was applied to identify differently-expressed genes (DEGs) (Love et al., 2014). The cutoff (adjusted P-value < 0.001) recommended for a small-sample RNA-seq experiment was used (Soneson and Delorenzi, 2013). Gene ontology (GO) terms enriched among DEGs were analyzed with BiNGO and visualized with Cytoscape (Maere et al., 2005).

### Chromatin States Analysis

We used ChromHMM (Ernst and Kellis, 2012) with default parameters to characterize the chromatin state maps for control and Pi deficiency samples. We used the published profiles of H2A.Z ChIP-seq (Zahraeifard et al., 2018) and nucleosome occupancy (Zhang et al., 2018)

(MNase-seq), as well as the H3K4me3 profile generated in this study. All input data were binarized with BinarizedBam, included in ChromHMM (Ernst and Kellis, 2012), and input genomic DNA was used to adjust binarization thresholds locally. The common model of chromatin states in both control and Pi-deficiency samples was developed by concatenating the marks using a hidden Markov model. Five chromatin states were generated based on the learned model parameters as described in ChromHMM (Ernst and Kellis, 2012). Chromatin state changes were analyzed using a previously described method (Fiziev et al., 2017). Briefly, the control and –Pi genomes were divided into 200-bp bins, each occupied by one chromatin state, and the chromatin state annotations of control and Pi-deficiency genomes were overlapped. The number of bins in each possible chromatin state were counted and called as the observed number. The expected number was calculated by multiplying the number of bins in the two chromatin states involved in each transition (a change in transition from control to Pi deficiency sample) and divided by total bins in the genome to calculate enrichment scores. Similarity between each pair of chromatin states was controlled by dividing the enrichment scores of each state transition to the enrichment scores of the reverse state transition. The distribution of chromatin states were identified using CEAS software (Shin et al., 2009). Each protein coding gene was assigned to one chromatin state based on the state of the 200-bp bin encompassing the transcription start site. For bootstraping analysis, we used a custom FORTRAN script (Zahraeifard et al., 2018) to obtain the same number of randomly selected genes and estimate the percentage of overlap between these genes and each group of state transitions (1000 iterations). Binomial distribution tests were carried out with R (pbinom, P-value < 1.00E-03). For the chromatin state transition plot (Figure 4), chromatin states in control samples were differentially color-coded. Genes in each control chromatin state were sorted based on their positions within each chromosome. Chromatin transitions for each gene were connected with lines of colors matching those used for control chromatin states. Genes in each chromatin transition were positioned according to their expression changes upon Pi deficiency treatment, with up-regulated genes on the top and down-regulated on the bottom. These transition connections were plotted with ggplot2 (Wickham, 2016). For the circos plot (Figure 3), each rice chromosome was partitioned into bins of 5kbp. Chromatin states were merged from 200bp bins to 5kbp bins in both control and Pi deficiency samples. The most dominant chromatin state in each merged bin, or the chromatin state of the previous bin if most dominant chromatin state cannot be determined, was selected as the chromatin state for that bin. For gene type partition, the most dominant gene type, in base pair, was used as the bin type for each bin. Chromatin states, differential expression status, bin types for the merged bins were determined using customized scripts and visualized with an R package circlize (Gu et al., 2014).

## Supplemental Material

**Supplemental Figure S1.** H3K4me3 abundance is strongly correlated with gene length but varies in abundance at the transcription start site (TSS). H3K4me3 heat map (A) and average plot (B) based on the gene length in gene body of protein-coding genes (PCG) from 500 bp upstream of the transcription start site (TSS) to 500 bp downstream of the transcription termination site (TES). Five quintiles were ordered by gene length (Q1-Q5). Average plot (C) and heat map (D) of k-means H3K4me3 clusters around the TSS (50 bp upstream and downstream of the TSS). Control input reads were used for ChIP-Seq read normalization.

**Supplemental Figure S2.** Difference of H3K4me3 enrichment pattern across rice protein coding genes (PCG) under 24-hours of Pi deficiency. Average plot of H3K4me3 for all PCG in control (Ctrl) or Pi deficiency (–Pi) samples.

**Supplemental Figure S3.** Log2 fold change of genomic bins occupied by each chromatin state in control (ctrl) compared to Pi deficiency (-Pi) samples.

**Supplemental Figure S4.** CS5-CS1 and CS4-CS5 transitions occur in sequence. Bootstrapping analysis showing the percentage of CS5-CS1 genes that exhibit a CS4-CS5 transition in the bin downstream (A) or upstream (C) of the TSS. Bootstrapping analysis showing the percentage of CS4-CS5 genes that exhibit a CS5-CS1 transition in the bin upstream (B) or downstream (D) of the TSS. All data are means (±SD) for 1000 iterations. Asterisks indicate significance at p-value < 0.001.

**Supplemental Figure S5.** Identification of differentially expressed genes in response to 24 hours of Pi deficiency. (A) MA plots of RNA-seq data by DESeq2. Networks representing Gene Ontology (GO) terms in the Biological Process (B), cellular component (C) and molecular functions (M) category enriched in DEGs that are down-regulated (B) and up-regulated (C) by Phosphate deficiency. Bingo and Cytoscape were used to identify and visualize enriched GO terms. Circle color shows p-value of enrichment.

**Supplemental Figure S6.** Bootstrapping analysis showing the overlap between genes exhibiting chromatin state transitions (CS5-CS1, CS4-CS5) and genes down-regulated in shoots following a 21-day Pi deficiency treatment from a previous study (Secco et al., 2013). All data are means (±SD) for 1000 iterations. Asterisks indicate significance at p-value < 0.01.

**Supplemental Figure S7.** Gene Ontology (GO) terms enriched in differentially expressed genes (DEGs) that have a chromatin state (CS) transition of CS4-CS3. Bingo and Cytoscape were used to identify and visualize enriched GO terms. Circle color shows p-value of enrichment.

**Supplemental Table S1.** Summary of ChIP-seq and RNA-seq libraries (short reads). The number of total and uniquely mapped reads for shoots from 36-day-old rice seedlings under control conditions (Ctrl) or following a 24-hour P-deficiency treatment (-P). Each sample contains two replicates (rep).

**Supplemental Table S2.** Summary of gene ontology (GO) analysis of genes differentially expressed under P-deficiency.

**Supplementary Dataset 1.** Significantly enriched GO terms for four sub-groups of protein-coding genes displaying different H3K4me3 abundance levels at the TSS.

**Supplementary Dataset 2.** Significantly enriched GO terms for eight gene groups exhibiting specific chromatin transitions.

**Supplementary Dataset 3.** Differentially expressed genes showing a CS4-CS3 chromatin transition.

## Accession numbers

H3K4me3 ChIP-seq and RNA-seq data sets from this article were submitted to the NCBI Sequence Read Archive (SRA) Database under the accession, SRP102668.

## Acknowledgements

The authors thank High Performance Computing at Louisiana State University (HPC@LSU) for providing computer resources. We also thank Aliasghar Sepehri for sharing the custom FORTRAN script to perform bootstrapping analyses.

