## supplemental for "Defining chromatin state transitions predicts a network that modulates cell wall remodeling in phosphate-starved rice shoots"

| | | Total reads | Uniquely mapped reads (%) |
|---|---|---|---|
| Ctrl | H3K4me3-rep1 | 41066109 | 78.5 |
| | H3K4me3-rep2 | 32132970 | 63.7 |
| | RNA-seq-rep1 | 58992471 | 67.57 |
| | RNA-seq-rep2 | 65082473 | 68.36 |
| -Pi | H3K4me3-rep1 | 37099052 | 77.9 |
| | H3K4me3-rep2 | 36468290 | 78.5 |
| | RNA-seq-rep1 | 64365952 | 68.17 |
| | RNA-seq-rep2 | 62848509 | 68.48 |

Supplemental Table S1. Summary of ChIP-seq and RNA-seq libraries (short reads). The number of total and uniquely mapped reads for shoots from 36-day-old rice seedlings under control conditions (Ctrl) or following a 24-hour P-deficiency treatment (-P). Each sample contains two replicates (rep).

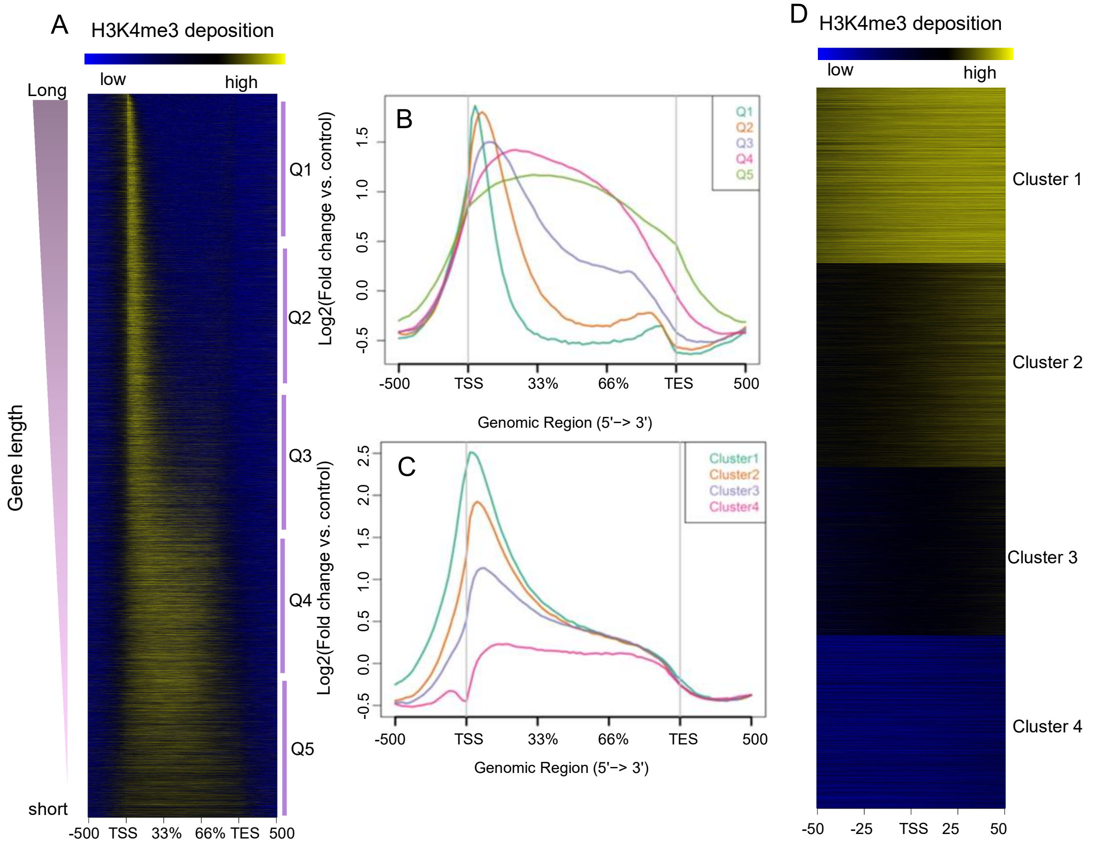


Supplemental Figure S1. H3K4me3 abundance is strongly correlated with gene length but varies in abundance at the transcription start site (TSS). H3K4me3 heat map (A) and average plot (B) based on the gene length in gene body of protein-coding genes (PCG) from 500 bp upstream of the transcription start site (TSS) to 500 bp downstream of the transcription termination site (TES). Five quintiles were ordered by gene length (Q1-Q5). Average plot (C) and heat map (D) of k-means H3K4me3 clusters around the TSS (50 bp upstream and downstream of the TSS). Control input reads were used for ChIP-Seq read normalization.

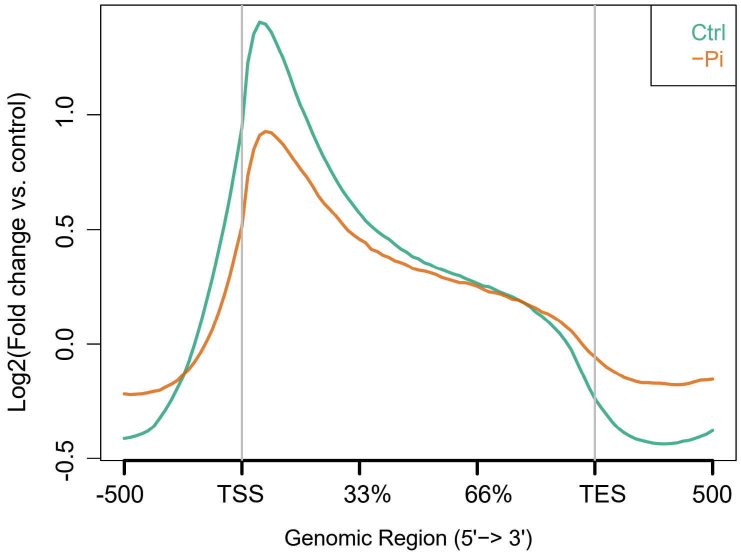


Supplemental Figure S2. Difference of H3K4me3 enrichment pattern across rice protein coding genes (PCG) under 24-hours of Pi deficiency. Average plot of H3K4me3 for all PCG in control (Ctrl) or Pi deficiency (–Pi ) samples.

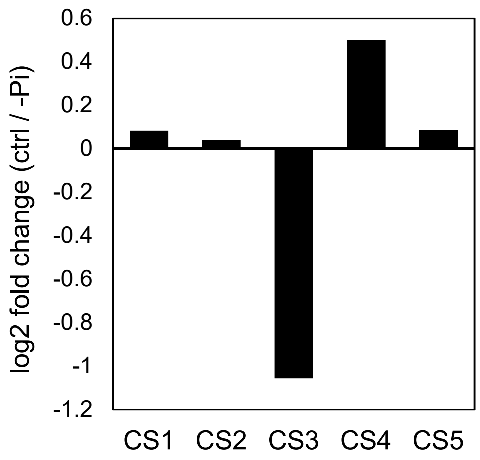


Supplemental Figure S3. Log2 fold change of genomic bins occupied by each chromatin state in control (ctrl) compared to Pi deficiency (-Pi) samples.

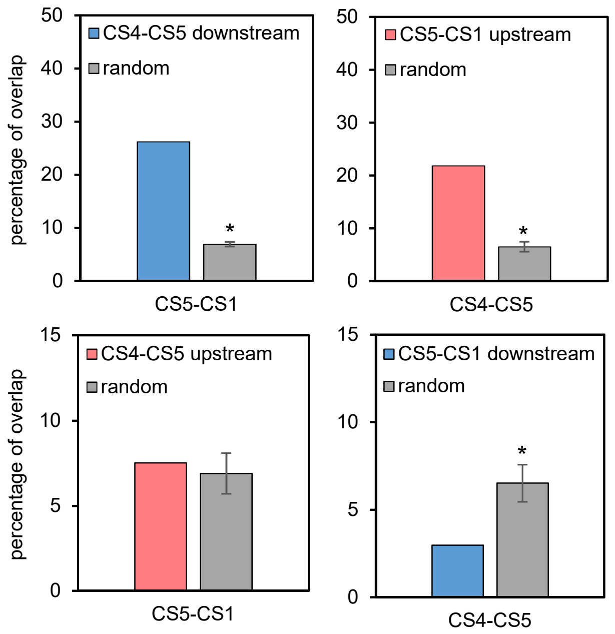


Supplemental Figure S4. CS5-CS1 and CS4-CS5 transitions occur in sequence. Bootstrapping analysis showing the percentage of CS5-CS1 genes that exhibit a CS4-CS5 transition in the bin downstream (A) or upstream (C) of the TSS. Bootstrapping analysis showing the percentage of CS4-CS5 genes that exhibit a CS5-CS1 transition in the bin upstream (B) or downstream (D) of the TSS. All data are means (±SD) for 1000 iterations. Asterisks indicate significance at p-value < 0.001.

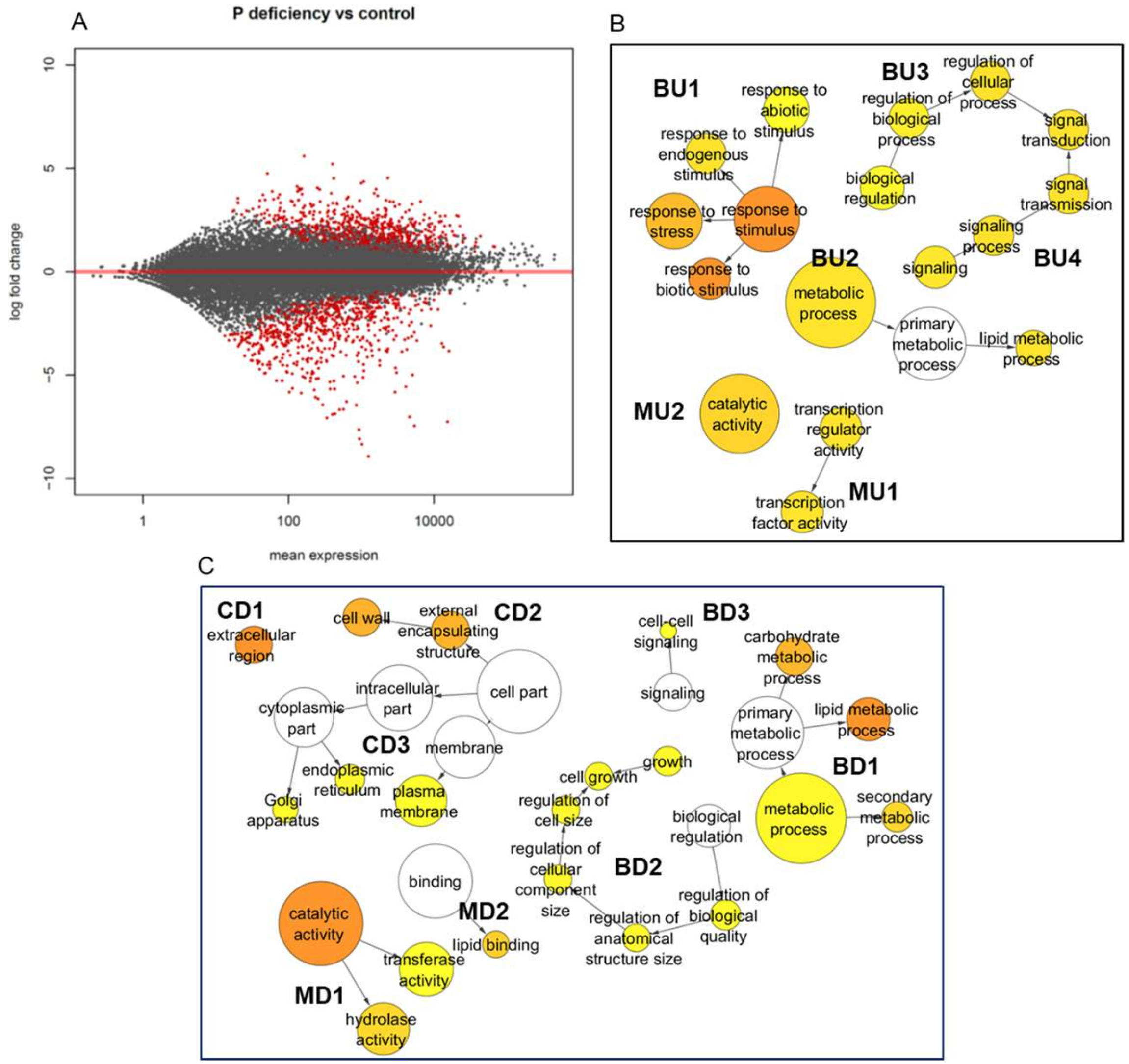


Supplemental Figure S5. Identification of differentially expressed genes in response to 24 hours of Pi deficiency. (A) MA plots of RNA-seq data by DESeq2. Networks representing Gene Ontology (GO) terms in the Biological Process (B), cellular component (C) and molecular functions (M) category enriched in DEGs that are down-regulated (B) and up-regulated (C) by Phosphate deficiency. Bingo and Cytoscape were used to identify and visualize enriched GO terms. Circle color shows p-value of enrichment.

| Category | -Pi response | Network | GO terms | Number (%) in DEGS | Number (%) in background | Log (P-value) |
|---|---|---|---|---|---|---|
| Biological process (B) | Up | BU1 | response to stimulus<br>-response to biotic stimulus<br>-response to stress<br>-response to endogenous stimulus<br>-response to abiotic stimulus | 186(30.7)<br>63(10.4)<br>130(21.5)<br>59(9.7)<br>79(13) | 5265(20.7)<br>1076(4.2)<br>3618(14.2)<br>1491(5.9)<br>2194(8.6) | -8.6<br>-10.3<br>-6.2<br>-4<br>-3.8 |
| | | BU2 | metabolic process<br>-lipid metabolic process | 381(62.9)<br>42(6.9) | 14183(55.7)<br>991(3.9) | -3.8<br>-3.6 |
| | | BU3 | regulation of biological process<br>-regulation of cellular process | 58(9.6)<br>58(9.6) | 1588(6.2)<br>1464(5.8) | -3.1<br>-4 |
| | | BU4 | signaling process<br>-signal transduction | 58(9.6)<br>58(9.6) | 1481(5.8)<br>1464(5.8) | -3.8<br>-4 |
| | Down | BD1 | metabolic process<br>-lipid metabolic process<br>-carbohydrate metabolic process<br>-secondary metabolic process | 383(61.5)<br>70(11.2)<br>50(8)<br>28(4.5) | 14183(55.7)<br>991(3.9)<br>972(3.8)<br>448(1.9) | -2.7<br>-14.8<br>-6.1<br>-4.5 |
| | | BD2 | growth<br>regulation of cell size | 26(4.2)<br>22(3.5) | 581(2.3)<br>465(1.8) | -2.6<br>-2.6 |
| | | BD3 | cell-cell signaling | 4(0.6) | 31(0.1) | -2.2 |
| Cellular component(C) | Down | CD1 | extracellular region | 47(7.5) | 559(2.2) | -12.7 |
| | | CD2 | external encapsulating structure<br>-cell wall | 50(8)<br>50(8) | 953(3.7)<br>943(3.7) | -6.4<br>-6.4 |
| | | CD3 | plasma membrane<br>Golgi apparatus<br>endoplasmic reticulum | 106(17)<br>17(2.7)<br>26(4.2) | 3116(12.2)<br>337(1.3)<br>562(2.2) | -3.6<br>-2.4<br>-2.8 |
| Molecular function (M) | Up | MU1 | transcription factor activity | 66(10.9) | 1773(7) | -3.7 |
| | | MU2 | catalytic activity | 287(47.4) | 9920(39) | -4.9 |
| | Down | MD1 | catalytic activity<br>-hydrolase activity<br>-transferase activity | 328(52.6)<br>110(17.7)<br>119(19.1) | 9920(39)<br>3112(12.2)<br>3913(15.4) | -11.8<br>-4.4<br>-2.2 |
| | | MD2 | lipid binding | 20(3.2) | 271(1.1) | -4.9 |

Supplemental Table S2: Summary of gene ontology (GO) analysis of genes differentially expressed under P-deficiency.

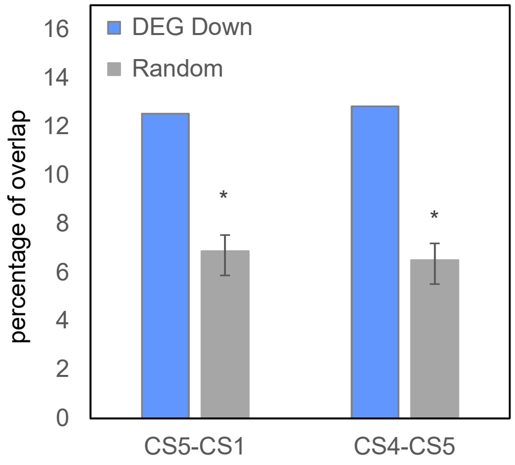


Supplemental Figure S6. Bootstrapping analysis showing the overlap between genes exhibiting chromatin state transitions (CS5-CS1, CS4-CS5) and genes down-regulated in shoots following a 21-day Pi deficiency treatment from a previous study (Secco et al., 2013). All data are means (±SD) for 1000 iterations. Asterisks indicate significance at p-value < 0.01.

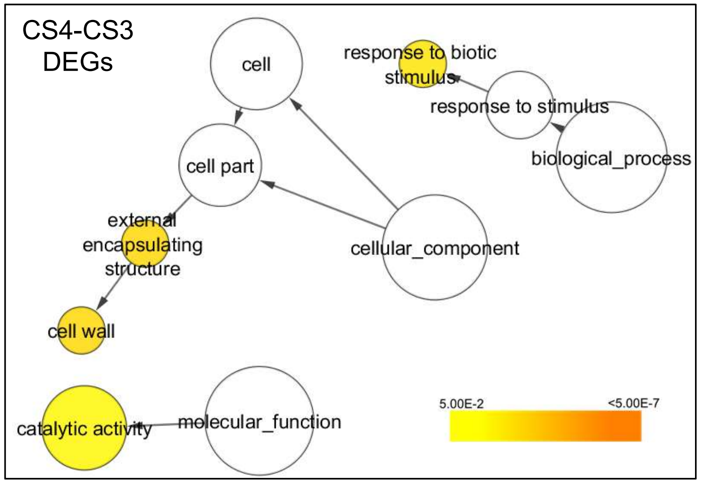


Supplemental Figure S7. Gene Ontology (GO) terms enriched in differentially expressed genes (DEGs) that have a chromatin state (CS) transition of CS4-CS3. Bingo and Cytoscape were used to identify and visualize enriched GO terms. Circle color shows p-value of enrichment.
